# Hippocampal sharp wave-ripple dynamics in NREM sleep encode motivation for anticipated physical activity

**DOI:** 10.1101/2023.03.14.532638

**Authors:** Chelsea M. Buhler, Julia C. Basso, Daniel Fine English

**Affiliations:** School of Neuroscience, Virginia Tech, VA 24061 USA; Department of Human Nutrition, Foods, and Exercise, Virginia Tech, VA, 24061 USA; Center for Health Behaviors Research, Fralin Biomedical Research Institute at VTC, VA, 24061 USA

**Keywords:** exercise, physical activity, voluntary wheel running, hippocampus, CA1, sharp wave-ripples, learning, memory, motivation, anticipation, reward, planning, incentive salience, metabolism, embodiment, sleep

## Abstract

Physical activity is an integral part of every mammal’s daily life, and as a driver of Darwinian fitness, required coordinated evolution of the body and brain. The decision to engage in physical activity is driven either by survival needs or by motivation for the rewarding qualities of physical activity itself. Rodents exhibit innate and learned motivation for voluntary wheel running, and over time run longer and farther, reflecting increased incentive salience and motivation for this consummatory behavior. Dynamic coordination of neural and somatic physiology are necessary to ensure the ability to perform behaviors that are motivationally variable. Hippocampal sharp wave-ripples (SWRs) have evolved both cognitive and metabolic functions, which in modern mammals may facilitate body-brain coordination. To determine if SWRs encode aspects of exercise motivation we monitored hippocampal CA1 SWRs and running behaviors in adult mice, while manipulating the incentive salience of the running experience. During non-REM (NREM) sleep, the duration of SWRs before (but not after) running positively correlated with future running duration, and larger pyramidal cell assemblies were activated in longer SWRs, suggesting that the CA1 network encodes exercise motivation at the level of neuronal spiking dynamics. Inter-Ripple-intervals (IRI) before but not after running were negatively correlated with running duration, reflecting more SWR bursting, which increases with learning. In contrast, SWR rates before and after running were positively correlated with running duration, potentially reflecting a tuning of metabolic demand for that day’s anticipated and actual energy expenditure rather than motivation. These results suggest a novel role for CA1 in exercise behaviors and specifically that cell assembly activity during SWRs encodes motivation for anticipated physical activity.

**SIGNIFICANCE STATEMENT:** Darwinian fitness is increased by body-brain coordination through internally generated motivation, though neural substrates are poorly understood. Specific hippocampal rhythms (i.e., CA1 SWRs), which have a well-established role in reward learning, action planning and memory consolidation, have also been shown to modulate systemic [glucose]. Using a mouse model of voluntary physical activity that requires body-brain coordination, we monitored SWR dynamics when animals were highly motivated and anticipated rewarding exercise (i.e., when body-brain coordination is of heightened importance). We found that during non-REM sleep before exercise, SWR dynamics (which reflect cognitive and metabolic functions) were correlated with future time spent exercising. This suggests that SWRs support cognitive and metabolic facets that motivate behavior by coordinating the body and brain.

## Introduction

Throughout mammalian evolution, physical activity has been a significant driver of Darwinian fitness, with the body and brain co-evolving, concomitantly increasing in complexity at the level of structure and function ^1^. As spatial navigation behaviors increased, the concurrent need to learn and remember the environment did as well, enhancing success in foraging and reproduction ^2^. Today however, physical inactivity in humans has become a global health crisis ^3^, with sedentary behaviors increasing the risk for a range of health issues including obesity, diabetes, heart disease, stroke, and cancer ^4^. Conversely, physical activity improves physical and mental health ^5^, making understanding exercise motivation a critical and timely scientific and societal issue.

The hippocampus is functionally positioned to play an important yet relatively unexplored role in exercise motivation, as it supports learning and memory ^6^ (including reward learning), planning of future actions ^7–11^ and although less acknowledged is involved in somatic physiology ^12–14^. Critical to all of these functions are CA1 sharp wave-ripples (SWRs) ^10^, during which specific populations (i.e., cell assemblies) of CA1 neurons become highly active ^8,9,15–17^, selected by computations involving interaction between excitation and inhibition ^10,18,19^, sending the final product of hippocampal computation to the rest of the brain ^10,20–22^. Interestingly, recent findings have shown that in addition to cognitive functions (largely involving hippocampal-neocortical dialogue), SWRs serve a specific somatic function: dynamically regulating peripheral glucose levels by communicating with the lateral septum ^14^. The differential translation of hippocampal SWR output by neocortex ^16,21–26^ and lateral septum ^14,27^ may thus enable multiplexing of cognitive and somatic functions. Furthermore, multiple hippocampal rhythms including SWR and theta are entrained to and driven by respiration ^28–31^, furthering the evidence of the important somatic/metabolic role of SWRs. Considering these dynamic roles of SWRs as modulators between the body and brain, we hypothesized that SWRs serve essential functions in behaviors that require enhanced body-brain coordination including physical activity.

The decision to engage in physical activity can be driven by survival needs (e.g., escaping danger) or by the motivation for the rewarding nature of physical activity itself (e.g., running exercise). Physical activity, in the form of voluntary (but not forced) wheel running, holds positive incentive salience for rodents ^32–34^. Critically, motivation for wheel running exercise can be quantified through the magnitude of the rebound running response after days of wheel deprivation ^35^. We thus investigated dynamics of 142,988 SWRs during NREM sleep in a rodent model exhibiting high levels of motivation for this behavior ^36^. We monitored hippocampal local field potentials (LFP) before, during, and after two hours of voluntary exercise; both on consecutive days and after days of wheel deprivation (deprivation increases incentive salience and thus running levels) ^32^. Differences in daily motivation to run were associated with changes in sleep architecture and SWR dynamics including rates and durations. These results suggest a previously unknown role for SWRs in the motivation for physical activity.

## Results

### Plastic motivation for voluntary wheel running in BL/6J x FVB/NJ hybrid mice chronically implanted with silicon probes

To identify relationships between voluntary physical activity and hippocampal network oscillations we monitored wheel running (Fig 1A) and hippocampal extracellular field potentials (Fig 1B) in freely behaving mice under the SRS protocol (Fig 1A). Hybrid mice ^36^ were chosen because they are a more ethologically relevant model compared to inbred strains. We analyzed data only from days after which the wheel running experience was no longer novel (i.e., day 3 and on) because novelty generally produces variability in behavior across animals, and specifically alters SWR dynamics ^8–10,15,16,37^ (also see Methods). Time spent running increased over days (Fig 1D; r =0.59, p=0.001), and deprivation resulted in a subsequent rebound response upon return of the wheel (Fig 1C; t (3) =-3.798, p=0.032), confirming mice found wheel running rewarding. We observed canonical relationships between behavioral states and hippocampal oscillations ^10,22,38^: theta oscillations were most prevalent in wake and REM sleep, whereas SWRs were most prevalent in NREM sleep. Importantly, wheel running bouts were accompanied by sustained theta oscillations, resulting in a positive correlation between theta power and time spent running. (Fig 1E; r = 0.68, p=0.001). The SRS protocol thus meets the need of our study in that mice chronically implanted with silicon probes exhibit normal exercise behaviors and hippocampal activity.

**Figure 1:**
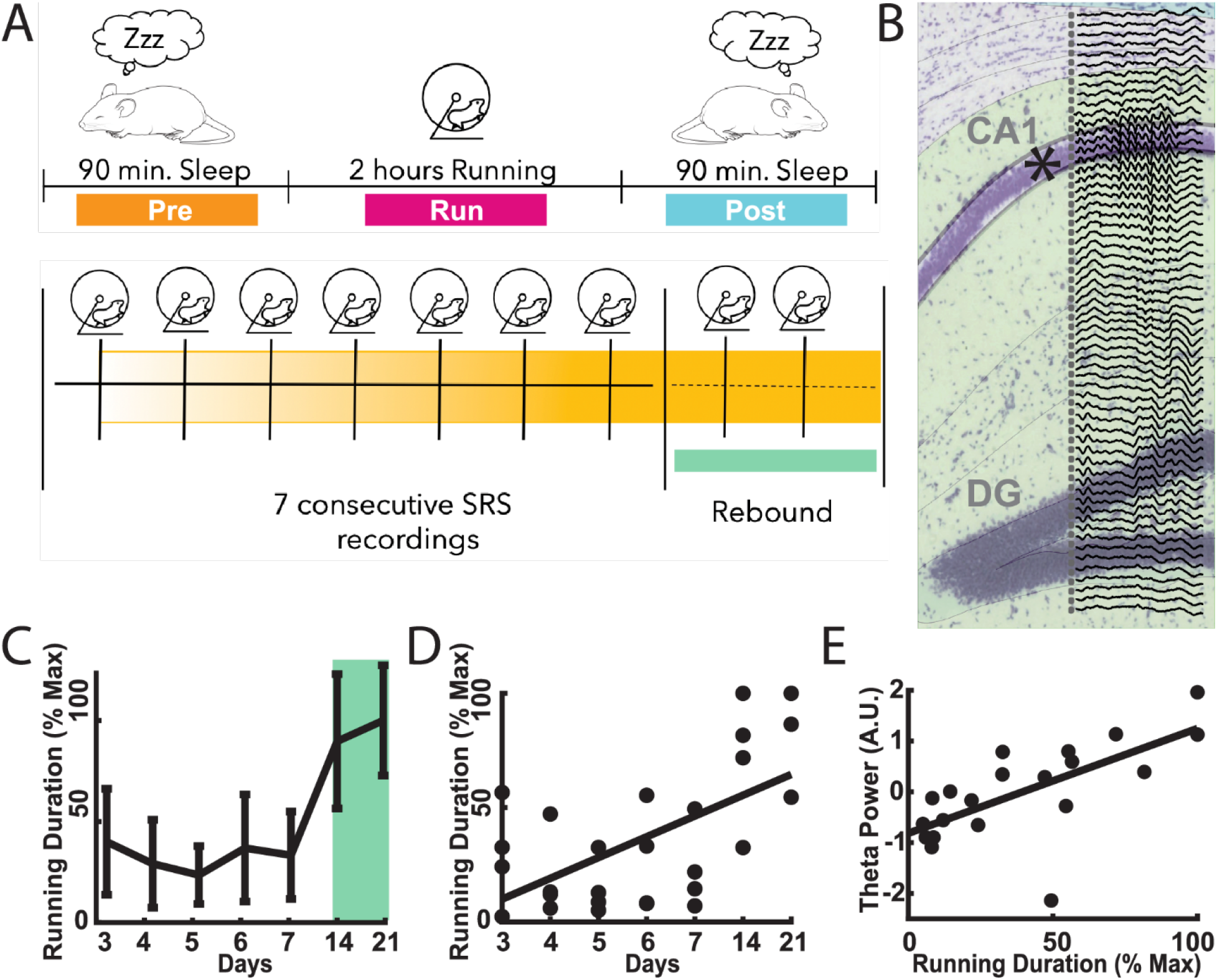
Motivation behaviors and hippocampal physiology in exercising mice. **A**. Behavioral Protocol. **B**. Example silicon probe recording CA1-DG 64 channel probe with 20 micron spacing implanted during SRS protocol. LFP traces are 150 ms illustrating a SWR event. **C**. Mean running duration on consecutive and rebound days. (t(3)= -3.798, p= 0.032) **D**. Running over Days (r= 0.59, p= 0.001); Pearson’s correlation. **E**. Hippocampal theta correlates with running duration (r= 0.68, p=0.001); Pearson’s correlation. **Each point is data from a single day, N=4 mice except on rebound 2 in which one mouse omitted because it was awake for all of Pre, resulting in 27 data points per correlation.

### Anticipation of rewarding exercise results in increased wakefulness and compressed NREM sleep

We quantified relative time spent in different sleep/wake states using established methods ^39^, Fig 2, and found that on rebound days mice spent significantly more time awake in Pre as compared to *Post* exercise epochs (p=0.01). We found that over days the percent time in NREM in Pre epochs was negatively correlated with future time spent running (Fig 2C; r=-.676, p=0.0001). While the opposite was true for NREM in *Post* epochs (Fig 2D; r=.426 p=0.026). Intriguingly, in Pre there is a significant negative correlation between SWR rates and the % time spent in NREM sleep (r=-0.6337, p= 0.0003), as well as significant negative correlations between inter-Ripple-interval (IRI) and future time spent running (Fig 3C, r=-0.50899, p=0.006), suggesting an anticipation driven compression of SWR mediated computations into a shorter duration of time. This was not the case for the *Post* epoch correlations between IRI and running duration (Fig 3C, r=-0.37591, p=0.0533) or SWR rates and the % time spent in NREM (r=0.363, p=0.06). Thus, periods of increased running were preceded, but not followed, by compressed NREM sleep.

**Figure 2:**
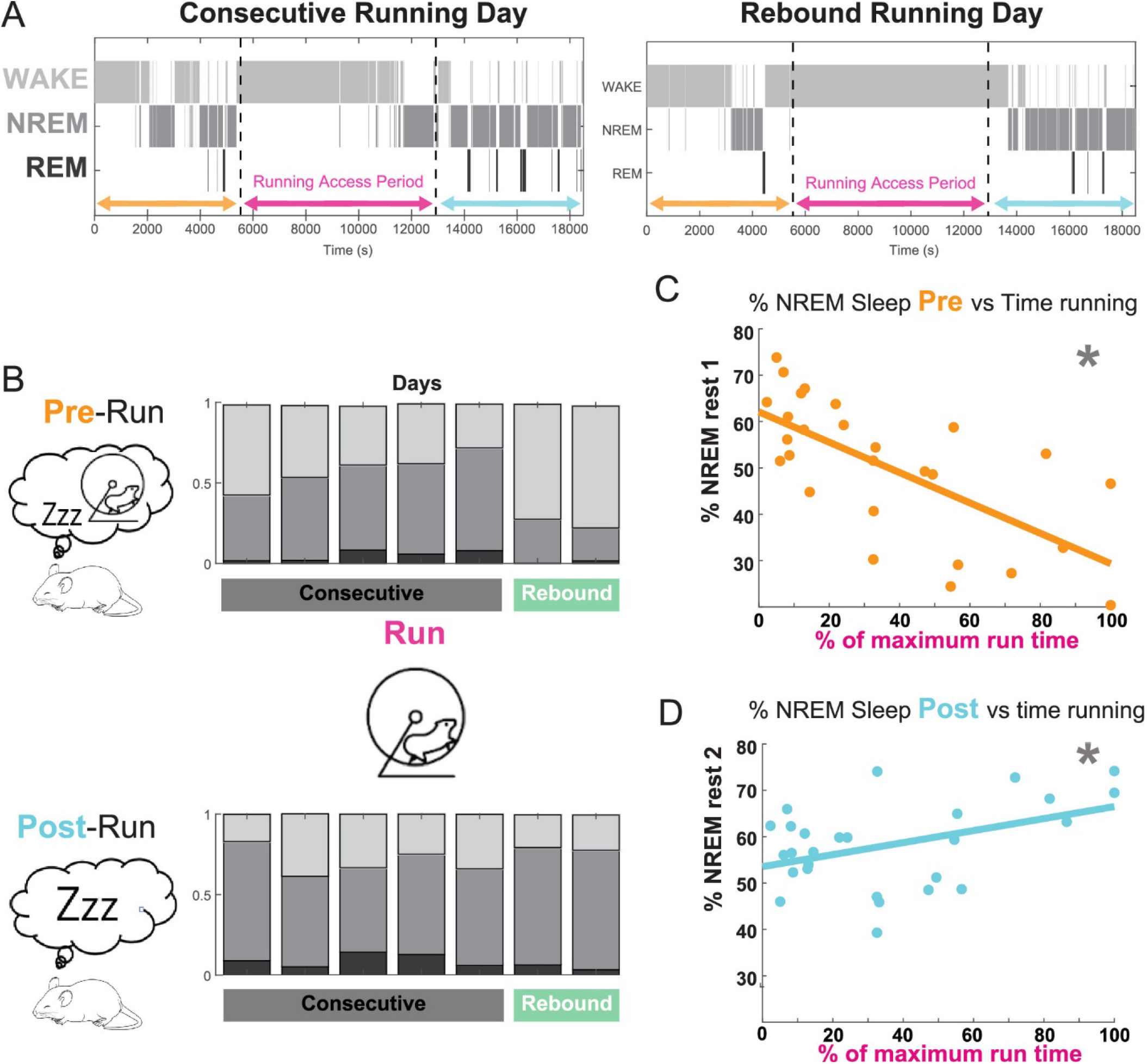
Alterations in wake/sleep dynamics associated with exercise motivation. **A**. Representative hypnograms of a consecutive day (left) and a rebound day (right), for the entire period of recording. **B**. Summary figure of the proportion of time spent in wake, NREM and REM states in Pre (top) and Post (bottom) on consecutive and rebound days for one mouse. **C**. Correlation between running and the percentage of time spent in NREM sleep during Pre (r=-.676, p=0.0001);Pearson’s correlation. **D**. Correlation between running and the percentage of time spent in NREM sleep during Post (r=.426 p=0.026);Pearson’s correlation. **Each point is data from a single day, N=4 mice except on rebound 2 in which one mouse omitted because it was awake for all of Pre, resulting in 27 data points per correlation.

**Figure 3:**
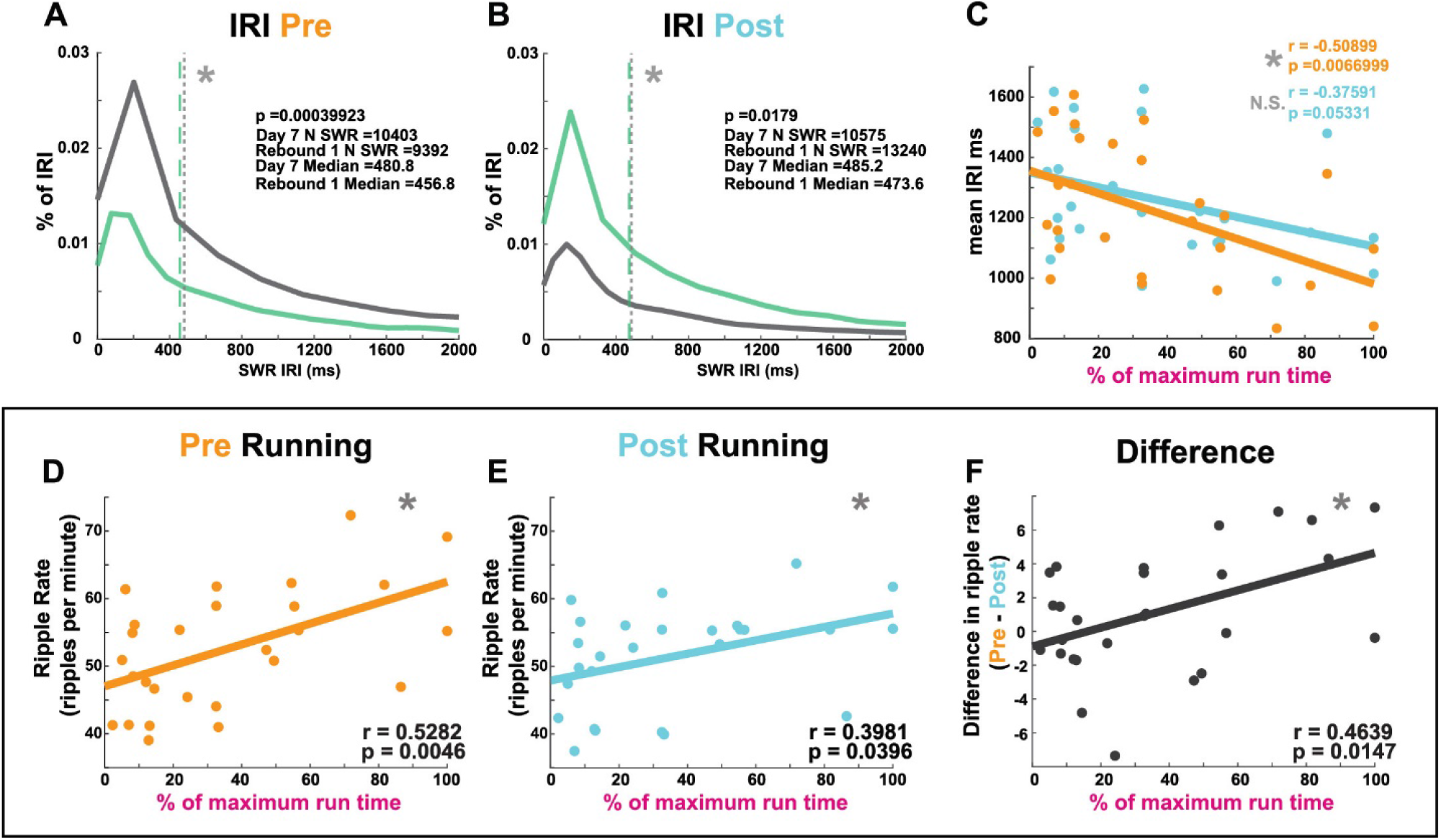
SWR Rate and inter-Ripple-interval correlate with daily physical activity. SWR rate (ripples per minute) in Pre and Post NREM sleep correlates with time spent running. **A** Kernel Distributions of IRI in Pre on day 7 (grey) and Rebound 1 (green) (n=10403 and 9392 SWR respectively, from 4 mice). Dashed lines indicate medians,(p<0.001);rank-sum test. **B** Kernel Distributions of IRI in Pre on day 7 (grey) and Rebound 1 (green) (n=10403 and 9392 SWR respectively, from 4 mice). Dashed lines indicate medians,(p=0.0178);rank-sum test. **C**. In Pre mean IRIs negatively correlate with running (r= -0.50899, p=0.0066999); Pearson’s correlation. In Post there is no significant correlation (r= -0.3759, p=0.05331); Pearson’s correlation. **D**. In Pre SWR rate significantly correlates with time spent running(r= 0.5282, p=0.0046); Pearson’s correlation. **E**. In Post SWR rate significantly correlates with time spent running(r= 0.3981, p=0.0396); Pearson’s correlation. **F**. The difference between SWR rate (Pre – Post) significantly correlates with time spent running (r= 0.4639, p=0.0147); Pearson’s correlation. **Each point is data from a single day, N=4 mice except on rebound 2 in which one mouse omitted because it was awake for all of Pre, resulting in 27 data points per correlation.

### SWR rates correlate with daily exercise performance

We compared SWRs in NREM sleep in *Pre* and *Post* run epochs in adult mice. Neither SWR mean power or frequency exhibited significant correlations with exercise levels (data not shown). Rates, IRI and durations of SWRs, however, were related to daily physical activity (Fig 3 & 4). The median duration of NREM IRIs are significantly lower on rebound 1 than Day 7 (Fig 3 A,B), despite the notable opposition in proportion of IRI in day 7 and rebound 1 from Pre to Post. Furthermore, NREM IRIs in Pre but not Post were positively correlated with time spent running (Fig 3C: Pre; r= -0.50899, p=0.0066999, Post; r= -0.3759, p=0.05331). NREM SWR rates in both *Pre* and *Post* epochs were positively correlated with daily running levels (Fig 3D: Pre; r= 0.5282, p=0.0046), (Fig 3E: Post; r= 0.3981, p=0.0396). The differences in SWR rate (Pre-Post) are also positively correlated with daily running performance (Fig 3F: Pre-Post; r= 0.4639, p=0.0147). Examined together these relationships between SWR rate/IRI and running performance indicate that the rate of SWRs may be related to the daily energy expenditure of the animals, supporting the metabolic/homeostatic drive both in preparation for physical exertion as well as in the recovery phase after physical exertion.

**Figure 4:**
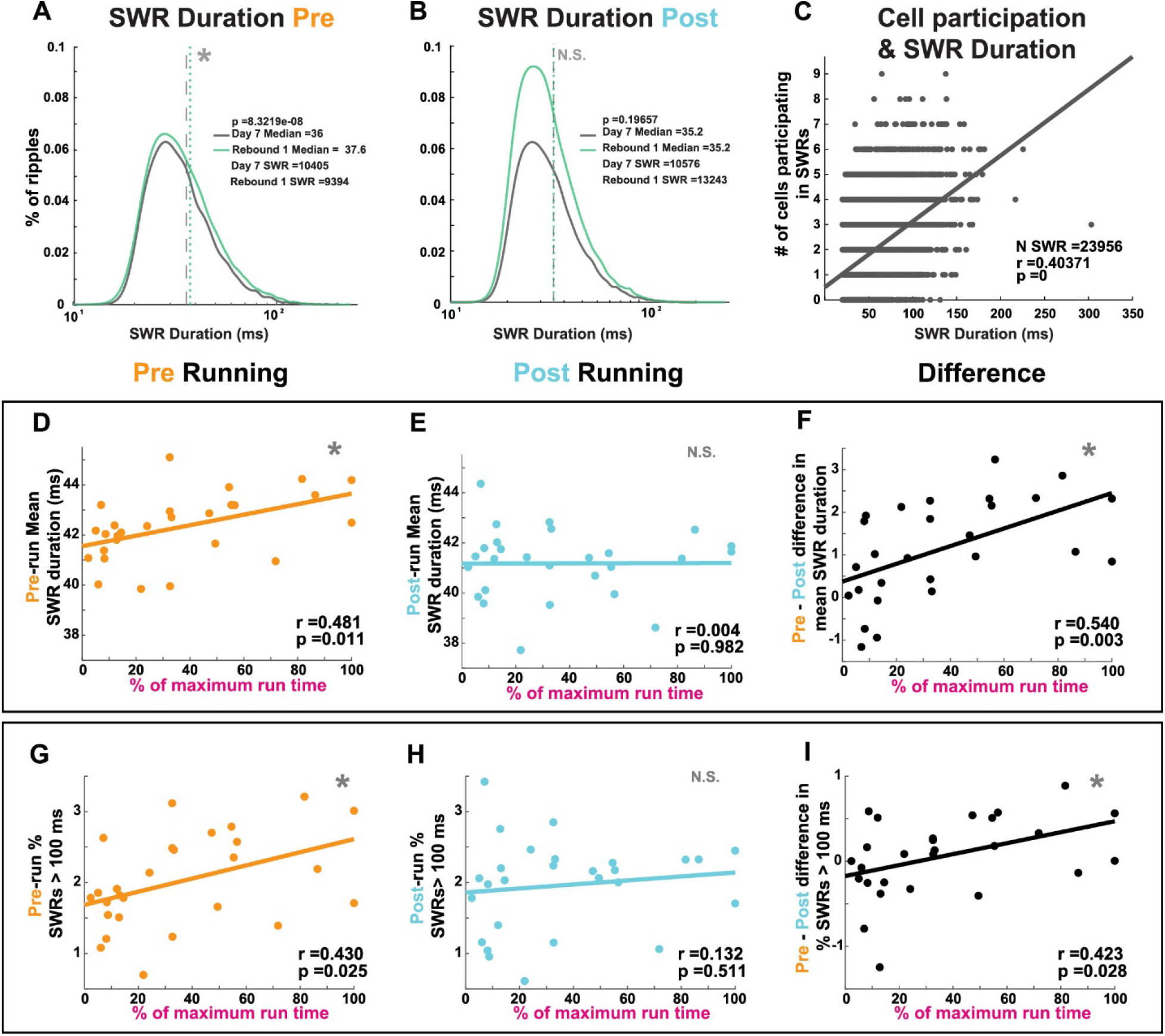
SWR Durations correlate with future physical activity. **A**. Kernel Distributions of SWR durations in Pre on day 7 (grey) and Rebound 1 (green) (n=10405 and 9394 SWR respectively, from 4 mice). Dashed lines indicate medians, p<0.001;rank-sum test. **B**. Kernel Distributions of SWR durations in Post on day 7 (grey) and Rebound 1 (green). (n=10576 and 13243 SWR respectively, from 4 mice). Dashed lines indicate medians, p=0.19657;rank-sum test. **C**. SWR duration significantly correlates with the number of CA1 Pyramidal cells spiking within SWR (n=23956 SWR, n=48 cells, from 4 recordings of 2 mice); r= 0.40371 p<0.001;Pearsons correlation. Sharp wave-ripple duration in Pre but not Post NREM sleep is correlated with time spent running. Relationships between Pre and Post run SWR duration **(D**,**E**,**F)** or percent of long duration SWRs **(G**,**H**,**I)** and time spent running. Note that SWR activity before but not after running, as well as the differences between these timepoints, are both significantly positively correlated with time spent running each day. **Each point is data from a single day, N=4 mice except on rebound 2 in which one mouse omitted because it was awake for all of Pre, resulting in 27 data points per correlation.

### SWR durations correlate with future exercise levels

Increased SWR duration is driven by the participation of more neurons (Fig 4C). SWR <u>durations</u> in *Pre* NREM were significantly positively correlated with future exercise levels which was not observed in *Post* NREM epochs (Fig 4: D, E, F). Furthermore, the distributions of SWR durations from day 7 and rebound 1 reveal significantly higher median SWR durations in Pre (Fig 4A: Pre; p< 0.001; rank-sum test) but not in Post (Fig 4B: Pre; p= 0.197; rank-sum test). In parallel, the percent of long duration ^40^ SWRs were significantly positively correlated with future exercise levels in *Pre* but not *Post* (Fig 4: G, H, I). These data suggest that SWRs dynamically encode the motivation for anticipated exercise.

To determine if the increased rate in SWRs was related to the observed changes in duration we calculated the correlation between these variables in the Pre running period, finding no significant relationship (r = -0.05, p = 0.81). This confirms that the SWR detection does not trivially reveal more long duration ripples as a function of SWR rate. More notably, these results indicate that distinct physiological mechanisms underlie changes in rate and duration of SWRs.

## Discussion

### Running is an innately rewarding consummatory behavior

Our data demonstrate that running is a rewarding behavior in mice. First, we demonstrated that mice increase their running over days. Second, we found that mice exhibit the rebound running response after deprivation, previously only shown in rats ^32,35^. Interestingly, this behavior was observed after only 7 days of acute running experience, while previous studies in rats reported that this behavior only occurs after habitual plateau levels of running are reached ^32,35^. Specifically, rebound running distance and rates on days 14 and 21 reached peak levels, above those demonstrated on the final consecutive day of running (day 7). This phenomenon is akin to what occurs after deprivation from other natural or pharmacological reinforcers such as food or drugs of abuse. To this point, we previously used a conditioned place preference model to show that running can be as rewarding as cocaine ^35^. These new findings demonstrate that motivation comes on board with only 7 days of consummatory behavior of wheel running experiences.

Furthermore, it is important to note that from the perspective of hippocampal rhythms, consummatory behaviors are typically associated with a cessation of theta oscillations and the occurrence of SWRs, however, exercise ^32–35^ and sex ^41,42^ are two prominent examples of innately rewarding consummatory behaviors which are associated not with SWRs but with sustained theta oscillations.

### Exercise modulates sleep architecture during anticipation of the running experience

When mice are anticipating the reward of running, they spend more time awake and less time asleep, with this effect being most prominent during rebound days to the point where some mice spend nearly their entire anticipatory period awake (Fig.2 A, B). Additionally, when mice fall asleep during this anticipatory period, they spend the majority of their time in NREM sleep with the percentage of NREM sleep negatively correlating to the time spent running (Fig 2C). That is, the more the mouse is motivated to run, the more time they spend awake and the less time they spend in NREM sleep before running. However, after the running experience, high levels of exercise motivation (i.e., time spent running) correspond to more NREM sleep (Fig. 2D). This latter finding is consistent with the human literature showing that both acute (a single bout) and chronic exercise produces longer periods of NREM sleep and shorter periods of REM sleep ^43^. Additionally, extant literature has revealed that exercise enhances overall quality of sleep including shorter sleep onset latency ^44^, time awake after sleep onset and longer total sleep time ^45^. We newly show that during an anticipatory running period, mice will regulate their sleep-wake cycles (Fig 2). That is, the external cue that the experience of running will soon be available alters the circadian pattern of the rodent such that awake periods are more prominent prior to (as compared to after) the running experience. Other literature has shown that such learned responses (e.g., food becoming available at a certain time of day) can alter circadian responses of other consummatory behaviors such as eating/feeding ^46^. Our findings reveal a general phenomenon regarding entrainment of innate behaviors (including physical activity and sleep), and that the timing of when these events occur in relation to one another can affect the temporal relationship of said behaviors.

### SWR dynamics encode exercise motivation and performance

SWRs are commonly studied using metrics including oscillation frequency, power, event duration, rate, and IRI. We found that SWR frequency and power were not related to exercise behaviors, as expected since SWR frequency is in general fixed across and during learning and memory tasks ^10^, with changes in frequency being associated with pathological states ^10,47,48^. In contrast, duration, rate, and IRI each demonstrated unique relationships with exercise motivation and performance which suggest a role for SWRs in multiple aspects of exercise behaviors. Of note, in rate, durations and IRI, all correlations were stronger for *Pre* as compared to *Post*, with the differences in rates (*Pre* - *Post*) showing positive correlation with exercise levels, suggesting potential anticipatory functions for both SWR rates, durations and IRI. The discrimination between rate and duration of SWRs is paramount, as distinct functions have been ascribed to modulations of each metric. Critically, we demonstrated that SWRs of longer duration recruit larger cell assemblies s (carrying more information). As such, dynamic SWR duration may enable flexible encoding of cognitive aspects of motivation, while SWR rate modulation may be more of an anticipatory and compensatory metabolic signal. In any case, these results suggest that the spiking content of SWRs is directly related to the anticipated and rewarding future exercise.

## Conclusions

We identified a novel role for SWRs in coordinating neuronal and physical activity: encoding motivation for future exercise in the activity of CA1 cell assemblies in SWRs. This suggests that both motor and cognitive processes are integrated within the same neuronal circuits - evidence for embodied cognition (i.e., the embodied brain). This perspective may help to reconcile the dichotomous theories of the hippocampus as a spatial map versus the seat of learning and memory. The hippocampus may thus be a core embodied center of the brain supporting somatic and cognitive processes, as well as their coordination through time. This perspective directly aligns with the co-evolution of the body and brain ^49^.

Importantly these results are novel and separate from SWR preplay ^50^ and replay ^8,9,15,16^. These concepts regard the structure of pre-existing and/or learned spatial maps. Our results instead introduce the novel concept that cell assembly dynamics in NREM SWRs encode a plastic representation of the motivation for a future innate consummatory behavior. We further hypothesize that incepting such activity could be used as a method to increase exercise motivation and performance.

## Materials and Methods

### Experimental considerations

For all experiments, we utilized four exercise naïve F1 BL/6J x FVB/NJ hybrid female mice bred in our colony. This mouse strain was utilized as previous work ^36^ has demonstrated high levels of cognitive functioning and low levels of anxiety compared to typical monogenic strains (e.g., C57), making them a more ethologically relevant model to study innate behaviors such as physical activity. Considering that this study focused on the motivation for voluntary exercise, we intentionally chose to use females, as the voluntary wheel running distances and rates are 1.5 times that of male, females run more consistently across the lifespan and have higher mean levels of physical activity ^32^, particularly at the adult timepoints we are studying. To limit inter-animal variability in both physiology and behavior, all animals were housed identically, and recordings and behavioral tasks were completed in the same behavioral testing rooms at the same time of day to insure that observed variabilities were not due to location or circadian rhythms. To minimize effects of age, young adult mice (∼PND 60) were used for this study, as wheel running is most robust during this time period. All mice received *ad libitum* food and water, as food and/or water deprivation have been shown to alter wheel running patterns^51^. Finally, room temperatures, light/dark cycles, noise level, and odors were monitored and kept constant to eliminate any additional stressors. All procedures were approved by the Institutional Animal Care and Use Committee at Virginia Tech.

### Behavioral task

We utilized the Sleep Run Sleep (SRS) protocol for measuring hippocampal activity and exercise behaviors. The SRS protocol was developed to measure hippocampal LFP biomarkers in the context of acute exercise experience. Under the SRS protocol (fig 1A), exercise-naïve female BL/6J x FVB/NJ mice were chronically implanted with a 64-channel silicon probe (Cambridge Neurotech) spanning the hippocampus from the CA1 through the dentate gyrus granular cell layer at 10–12 weeks old. Following surgical recovery, baseline recordings were made in the home cage to habituate mice to the recording procedures. Under the SRS protocol, neural activity is recorded throughout behavioral tests and sleep sessions; this protocol therefore enabled us to examine sleep architecture and signaling before and after behavior. These recordings were obtained during the animal’s light cycle to ensure the capture of substantive sleep physiology. The SRS protocol begins with 90 minutes of sleep in the home cage, followed by 2 hours of running wheel access in a novel cage, and ending with another 90 minutes of sleep in the home cage. The LFP was recorded continuously and synchronized with video of the running phase activity. Following the collection of baseline sleep activity, we collected 7 days of consecutive SRS recordings to examine effects across time. Due to neophobia, we found that between animals there was variability in interactions with the wheel on the first two days that stabilized by day 3. Therefore, we only considered data from day 3 and after in our analyses. Two rebound-SRS recordings were collected 7 and 14 days after the last consecutive day to examine rebound running behaviors and hippocampal activity during different motivational states. Running behaviors were analyzed using the MedAssociates, Inc. low-profile wireless running wheel for mice (ENV-047) and wheel running software, as well as manual scoring for total duration of running (seconds) per running phase. Overall, the SRS protocol (informed by our previous work on exercise motivation) allowed us to examine hippocampal physiology during both the initial phase of consecutive-day running and the rebound running response phase on days after deprivation ^35^.

### Chronic Hippocampal Extracellular recordings in freely behaving mice

Mice (N=4) were chronically implanted with 64-channel linear silicon probes (Cambridge NeuroTech) across the hippocampus, a described in (English et al, 2017). Briefly, probes were mounted on custom microdrives secured to the skull with dental cement, and subsequently inserted into the brain through a single craniotomy located at –1.75 AP and 1 ML in reference to bregma. A 50 μm stainless steel ground wire was implanted between the skull and dura over the cerebellum. This configuration allowed us to record electrical activity over >1 mm with 20-micron resolution and identify anatomically localized activity across the dentate gyrus (DG) and CA regions. Recordings were obtained using a digital multiplexing amplifier (Intan Technologies LLC) sampled at 30 kHz and synchronized with overhead video.

### Figure acknowledgements

Figure 1B was made with images from the Allen mouse brain atlas. Figures 1 and 2 contain images from BioRender.com.

### Analysis of Local Field Potentials

Analysis was performed using custom MATLAB scripts including those based on freely available functions from the Buzcode toolbox https://github.com/buzsakilab/buzcode.

### Theta power

Mean theta (7-11 Hz) power is computed using freely available functions from the FMAtoolbox (MTSpectrogram and SpectrogramBands).

### Ripple Detection

Ripple Detection was performed using standard algorithms as we have previously published ^18,52^.

### State Scoring

Segmentation of data into brain states (i.e., Wake, REM, NREM) was performed using previous methods ^39^. Briefly, periods of each state were segregated based upon a combination of characteristic features of LFP and EMG activity, followed by manual curation using accelerometer data to validate state scoring.

### Statistics

All statistical analyses were performed in MATLAB (MathWorks). We observed highly variable behavior on the first and/or second days in different animals (data not shown), which were no longer present by day 3, as such statistical analyses are restricted to day 3 forward with the exclusion of days in which sleep was not captured within resting epochs. Considering that significant inter-individual variability in running distance/duration was observed, we normalized wheel running duration within animals (% of max running duration = daily duration/maximum daily duration), and cross-validated normalization procedures to make accurate and robust quantification of exercise. Note, the maximum daily duration that was used to normalize running behaviors was taken across all experimental days to establish an individual exercise performance scale. We investigate the relationship of daily wheel running behaviors to all variables of interest (e.g., SWR rate and duration) by computing the Pearson’s linear correlation coefficient and p-values using a student’s t distribution for a transformation of the correlation. All correlations contain 27 data points from N=4 mice (one day is omitted because there was no NREM sleep in Pre).

## ACKNOWLEDGMENTS

DFE is supported by grants from The Simons Foundation and The Whitehall Foundation. JCB is supported by the iTHRIV Scholars Program, which is supported in part by the National Center for Advancing Translational Sciences of the NIH (UL1TR003015 *and* KL2TR003016). CMB is supported by the Virginia Tech School of Neuroscience.

## Supplementary Figures

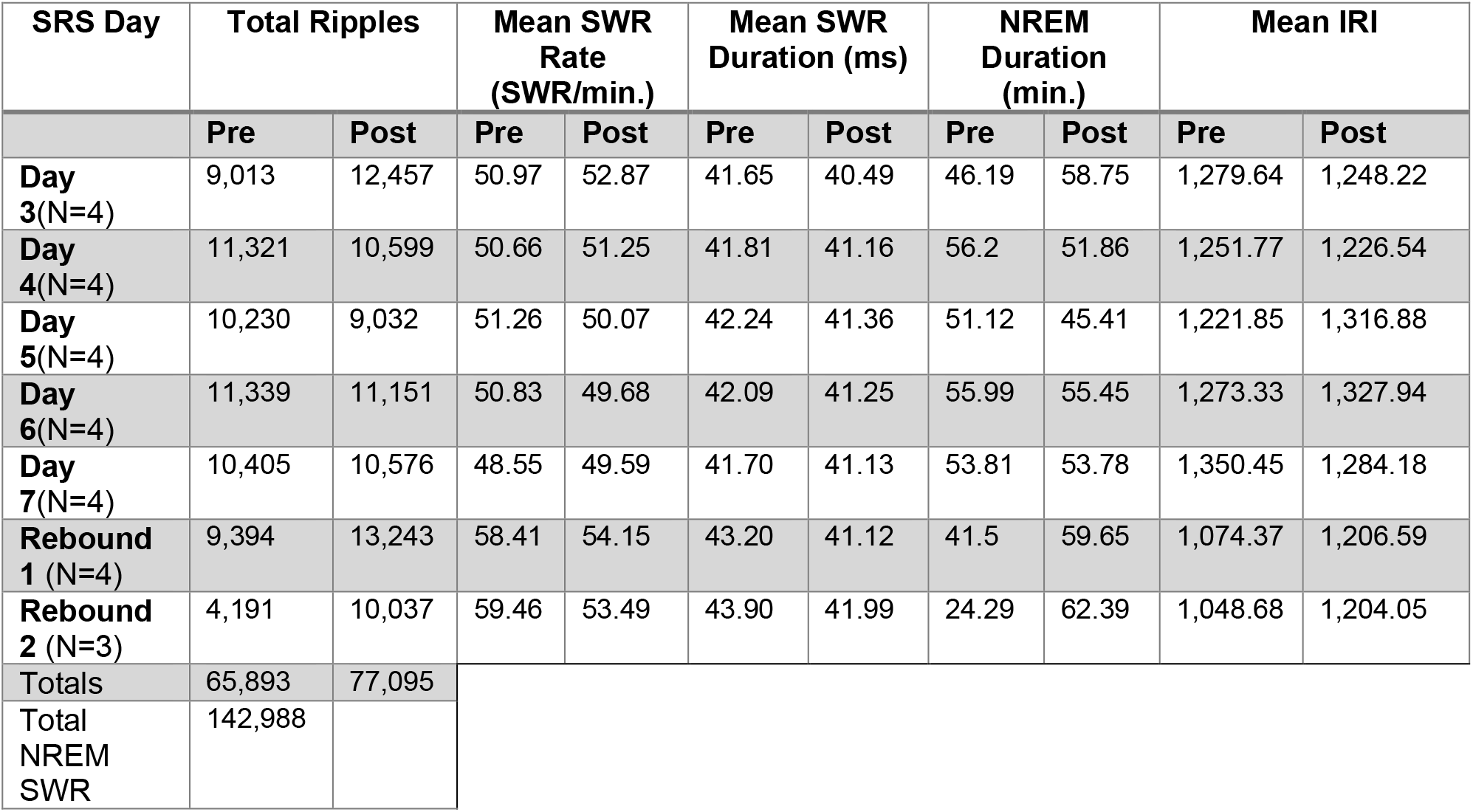

